# Protection against *Clostridioides difficile* disease by a naturally avirulent *C. difficile* strain

**DOI:** 10.1101/2024.05.06.592814

**Authors:** Qiwen Dong, Stephen Harper, Emma McSpadden, Sophie S. Son, Marie-Maude Allen, Huaiying Lin, Rita C. Smith, Carolyn Metcalfe, Victoria Burgo, Che Woodson, Anitha Sundararajan, Amber Rose, Mary McMillin, David Moran, Jessica Little, Michael Mullowney, Ashley M. Sidebottom, Aimee Shen, Louis-Charles Fortier, Eric G. Pamer

## Abstract

*Clostridioides difficile (C. difficile)* strains belonging to the epidemic BI/NAP1/027 (RT027) group have been associated with increased transmissibility and disease severity. In addition to the major toxin A and toxin B virulence factors, RT027 strains also encode the CDT binary toxin. Our lab previously identified a toxigenic RT027 isolate, ST1-75, that is avirulent in mice despite densely colonizing the colon. Here, we show that co-infecting mice with the avirulent ST1-75 and virulent R20291 strains protects mice from colitis due to rapid clearance of the virulent strain and persistence of the avirulent strain. Although avirulence of ST1-75 is due to a mutation in the *cdtR* gene, which encodes a response regulator that modulates the production of all three *C. difficile* toxins, the ability of ST1-75 to protect against acute colitis is not directly attributable to the *cdtR* mutation. Metabolomic analyses indicate that the ST1-75 strain depletes amino acids more rapidly than the R20291 strain and supplementation with amino acids ablates ST1-75’s competitive advantage, suggesting that the ST1-75 strain limits the growth of virulent R20291 bacteria by amino acid depletion. Since the germination kinetics and sensitivity to the co-germinant glycine are similar for the ST1-75 and R20291 strains, our results identify the rapidity of *in vivo* nutrient depletion as a mechanism providing strain-specific, virulence-independent competitive advantages to different BI/NAP1/027 strains. They also suggest that the ST1-75 strain may, as a biotherapeutic agent, enhance resistance to CDI in high-risk patients.

**Importance:** *Clostridioides difficile* infections (CDI) are prevalent in healthcare settings and are associated with high recurrence rates. Therapies to prevent CDI, including recent FDA-approved live biotherapeutic products, are costly and have not been used to prevent primary infections. While a nontoxigenic *C. difficile* strain (NTCD-M3) protects against virulent CDI in animals and reduced CDI recurrence in a phase 2 clinical trial, protection against CDI recurrence in humans was variable and required high doses of the nontoxigenic strain. Here we show that an avirulent *C. difficile* isolate, ST1-75, efficiently outcompetes virulent *C. difficile* strains in mice when co-infected at a 1:1 ratio. Our data suggest that inter-strain competition results from ST1-75’s more rapid depletion of amino acids than the virulent R20291 strain. Our study identifies inter-strain nutrient depletion as a potentially exploitable mechanism to reduce the incidence of CDI.

## Introduction

*Clostridioides difficile* (*C. difficile*) is a Gram-positive, spore-forming anaerobe that is the leading cause of nosocomial infection in U.S. adults. It is estimated that *C. difficile* causes 223,900 cases and 12,800 deaths yearly with annual healthcare costs in excess of $1 billion dollars (1). *C. difficile* infection (CDI) symptoms range from asymptomatic colonization to mild diarrhea to severe pseudomembranous colitis, which can lead to mortality (2). Susceptibility to CDIs is associated with gut dysbiosis and the loss of microbiota-mediated colonization resistance (3). Thus, prior antibiotic treatment is the major risk factor for development of CDI. Antibiotic administration, however, remains the standard-of-care for treating *C. difficile* infections, even though this treatment perpetuates gut dysbiosis and contributes to a patient’s roughly 30% risk of CDI relapse (4).

Multiple strategies have been applied to prevent primary and recurrent CDI. Hospitals mitigate *C. difficile* transmission by implementing contact precautions, hand hygiene and environmental decontamination (5). However, the effectiveness of these intervention strategies varies from study to study and greatly depends upon healthcare workers’ compliance (6–10). Notably, optimizing antibiotic choice, timing, and duration of administration via antibiotic stewardship programs has reduced CDI rates by 24-60% (11–15). While vaccination confers long-lived protection against a range of pathogens, vaccines against *C. difficile* recently failed to show efficacy in protecting primary CDI in phase 3 trials (16, 17). Fecal microbiota transplantation (FMT) is the most effective strategy for treating and preventing recurrent CDI, and there are new FDA-approved live biotherapeutic products that reduce the incidence of recurrent CDI (18, 19).

Given FMT’s success, there is considerable interest in developing probiotic therapies that are less expensive and simpler to produce on a large-scale. Indeed, several studies have shown that probiotics can work as an adjuvant therapy to reduce the risk of CDI (20) by restoring the gut community and metabolites that restrict *C. difficile* growth. However, due to the complexity of the probiotic composition, administration timing, and duration of the studies, variable outcomes have been reported (21). Another bacteriotherapeutic strategy being explored is the use of nontoxigenic *C. difficile* strains to prevent recurrent CDI. One of such nontoxigenic *C. difficile* strains, NTCD-M3, provided protection against virulent CDI in hamsters and later showed efficacy in a phase 2b clinical study for preventing CDI relapse, with NTCD-M3 treatment being associated with a significantly reduced recurrence rate (11%) compared to the placebo group (30%) (22). Thus, identifying avirulent *C. difficile* strains that can serve as biotherapeutics for combatting virulent infection is of considerable interest (23–26). More complete mechanistic understanding of how avirulent *C. difficile* strains can prevent disease caused by virulent strains may provide additional approaches to prevent CDI.

We previously characterized a ribotype 027 *C. difficile* isolate, ST1-75, that is avirulent in mouse models despite encoding all three toxins, including *C. difficile*’s primary virulence factors Toxin A (TcdA), Toxin B (TcdB) and the binary toxin CDT. We showed that avirulence of ST1-75 is attributable to a natural mutation occurring in the *cdtR* gene, which encodes a response regulator for the binary toxin (27, 28). While CdtR regulates the expression of the gene encoding CDT, the natural in-frame deletion of 69 bp from the *cdtR* gene is sufficient to reduce the expression of not only the genes encoding CDT but also Toxin A and Toxin B (27, 29). Here, we found that coinfecting mice with the avirulent strain ST1-75 and the virulent strain R20291 prevents colitis induced by the virulent R20291 strain in mice and results in its rapid clearance from the gastrointestinal tract. Notably, protection against acute colitis by ST1-75 is not due to the mutation it harbors in its *cdtR* gene. Instead, metabolomic analyses indicate that ST1-75 more rapidly depletes amino acids, and spiking amino acids into nutrient-scarce media reduced ST1-75’s competitive advantage. Intestinal colonization with ST1-75 prior to antibiotic treatment reduced the risk of CDI following challenge with virulent R20291, suggesting that it may have potential as a prophylactic live biotherapeutic product.

## Results

### Avirulent ST1-75 protects mice from R20291-induced CDI colitis

Nontoxigenic or low virulence bacterial strains can provide protection against colitis induced by highly virulent strains. We tested whether the avirulent ST1-75 strain can prevent the virulent RT027 R20291 strain from causing disease when mice are co-infected with both strains. To sensitize mice to *C. difficile* infection, the mice were treated with an antibiotic cocktail consisting of metronidazole, neomycin, and vancomycin (MNV), in their drinking water for 3 days. Clindamycin was then administered by intraperitoneal injection 2 days after the MNV cocktail was stopped. Twenty-four hours later, mice were inoculated with either 200 *C. difficile* spores for the individual strains or 100 ST1-75 and 100 R20291 spores each during the co-infection (1:1 ratio, **Fig 1A**). As expected, mice infected with the avirulent ST1-75 strain alone did not exhibit weight loss or disease symptoms (based on an acute disease score), and the virulent R20291 strain induced severe weight loss and diarrhea (27). However, mice coinfected with both ST1-75 and R20291 did not exhibit any disease symptoms, similar to the ST1-75 infection alone (**Fig 1B-1C**), indicating that ST1-75 was able to protect mice from R20291-induced colitis. While comparable total fecal CFUs were measured for *C. difficile* for all three groups (**Fig 1D**), qPCR analyses revealed that R20291 comprised the minority of the fecal CFUs recovered from the co-infected mice (**Fig 1E**), suggesting that ST1-75 efficiently outcompetes R20291 *in vivo*. This intraspecies competition presumably allows ST1-75 to protect mice from developing *C. difficile-*induced colitis during coinfection.

**Figure 1.**
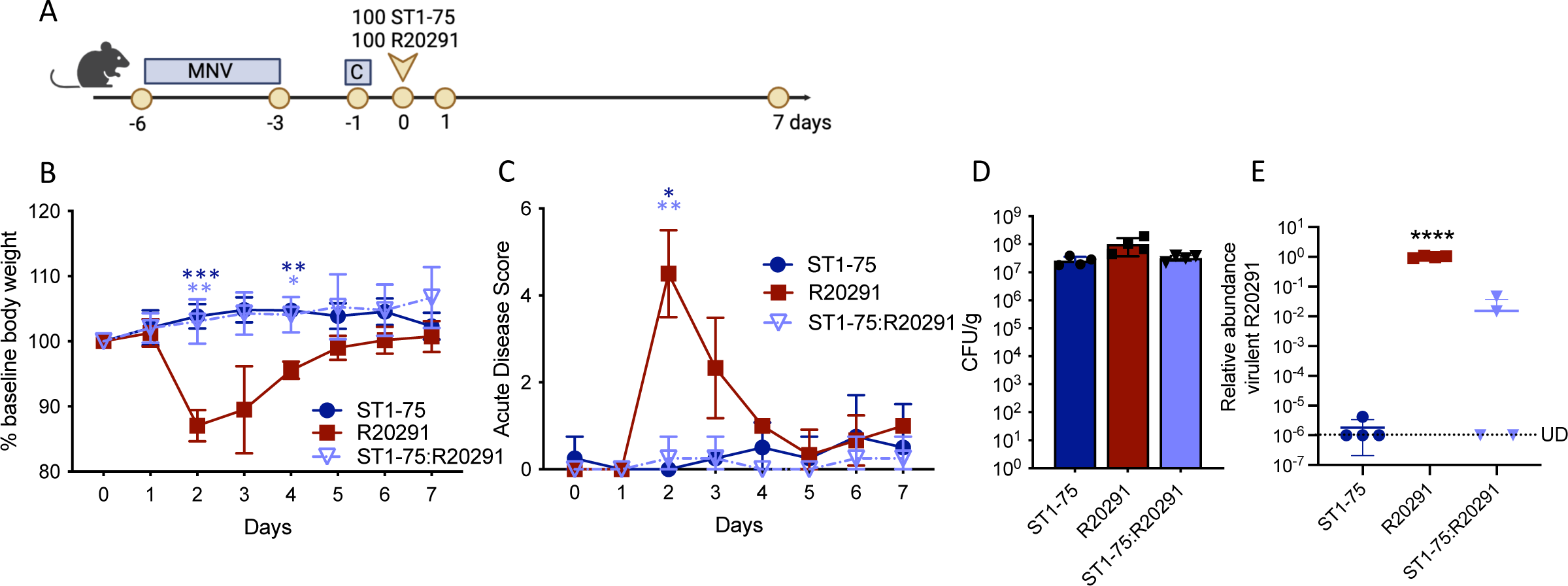
Coinfection of avirulent ST1-75 with a virulent *C. difficile* strain prevents disease in antibiotic-treated mice. (A) Schematic of the experimental procedure. Wild-type C57BL/6 mice (n = 4 per group) were treated with Metronidazole, Neomycin and Vancomycin (MNV, 0.25 g/L for each) in drinking water for 3 days, followed by one intraperitoneal injection of Clindamycin (200 mg/mouse), indicated as C in the schematic, 2 days after antibiotic recess. Then, mice were inoculated with 100 R20291 *C. difficile* spores and 100 ST1-75 *C. difficile* spores via oral gavage. Daily body weight and acute disease scores were monitored for 7 days post-infection. (B) %Weight loss relative to the baseline of mice infected with indicated strains. (C) Acute disease scores comprising weight loss, body temperature drop, diarrhea, and morbidity of mice infected with indicated strains. (D) Fecal colony-forming units were measured by plating on selective agar on 1 day post-infection. (E) Relative abundance of R20291 in feces from infected mice was determined by measuring wildtype *cdtR* copies by qPCR on 1 day post infection. UD: Under the limit of detection. Statistical significance was calculated by unpaired t-test and One-way ANOVA, * p < 0.05, ** p < 0.01, *** p < 0.001, **** p < 0.0001. Statistical significance was observed between R20291 to ST1-75 and R20291 to ST1-75:R20291.

To determine if there is a window of time in which ST1-75 can outcompete R20291 during infection, we assessed the ability of ST1-75 to prevent R20291-mediated colitis if mice were first challenged with R20291 and then ST1-75 was administered 6 hours afterwards at either a 1:1 ratio (200 R20291 and 200 ST1-75 spores) or a 1:50 ratio (200 R20291 and 10,000 ST1-75 spores). ST1-75 failed to prevent weight loss induced by R20291 in mice, even when ST1-75 was given in large (50-fold) excess (**Fig S1B**). Since this higher dose of ST1-75 administered 6 hours post-R20291 infection (blue diamonds) allowed ST1-75 to outcompete R20291 within 24 hrs at levels similar to the simultaneous 1:1 coinfection (light blue triangles), our results suggest that there is a critical window of time in which ST1-75 can prevent *C. difficile* pathogenicity.

Notably, even though R20291 predominated at the 24 hr timepoint when equal amounts of ST1-75 spores were administered 6 hours post-R20291 infection, ST1-75 eventually out-competed R20291 by Day 7 (green upside-down triangles) (**Fig S1C**). In contrast, when high doses of ST1-75 relative to R20291 were administered 24 hours post-R20291 infection, ST1-75 failed to outcompete R20291 even 7 days post-infection (maroon open circles) (**Fig S1C**). This result suggests that *C. difficile* establishes colonization resistance to secondary invading strains within 24 hours after infection. Taken together, our data reveal a competitive dynamic between ST1-75 and R20291 and indicate that ST1-75 retains its competitive fitness advantage even if administered a few hours after R20291.

### Protection by the avirulent ST1-75 is not due to differences in the lysogenic phages between strains

ST1-75 harbors two unique prophages in its genome, namely phiCD75-2 and phiCD75-3, which can become infectious upon induction (27). Prophages can determine the susceptibility of the bacterial host to phage infections via superinfection exclusion or other anti-phage defense systems (30–32). For example, ST1-75 is immune to infection and lysis induced by either phiCD75-2 or phiCD75-3 because ST1-75 carries these two prophages in its genome. In contrast, these same phages can infect and induce lysis in R20291. To assess whether prophages alter the competitive index between ST1-75 and R20291, we coinfected mice with ST1-75 and R20291 strains previously lysogenized with either phiCD75-2 (R_phiCD75-2) or phiCD75-3 (R_phiCD75-3), or both (R_ phiCD75-2/3) (**Fig 2A** and **Fig S2A**). Notably, lysogenized R20291 strains retained the ability to cause disease in mice (27) (**Fig 2B** and **Fig S2B**), but they were outcompeted by ST1-75 with similar kinetics to their parental R20291 strain (**Fig 2B-2D** and **Fig S2B-S2D**). Taken together, our results suggest that prophages within ST1-75 contribute minimally to the competitive advantage of ST1-75 over R20291.

**Figure 2.**
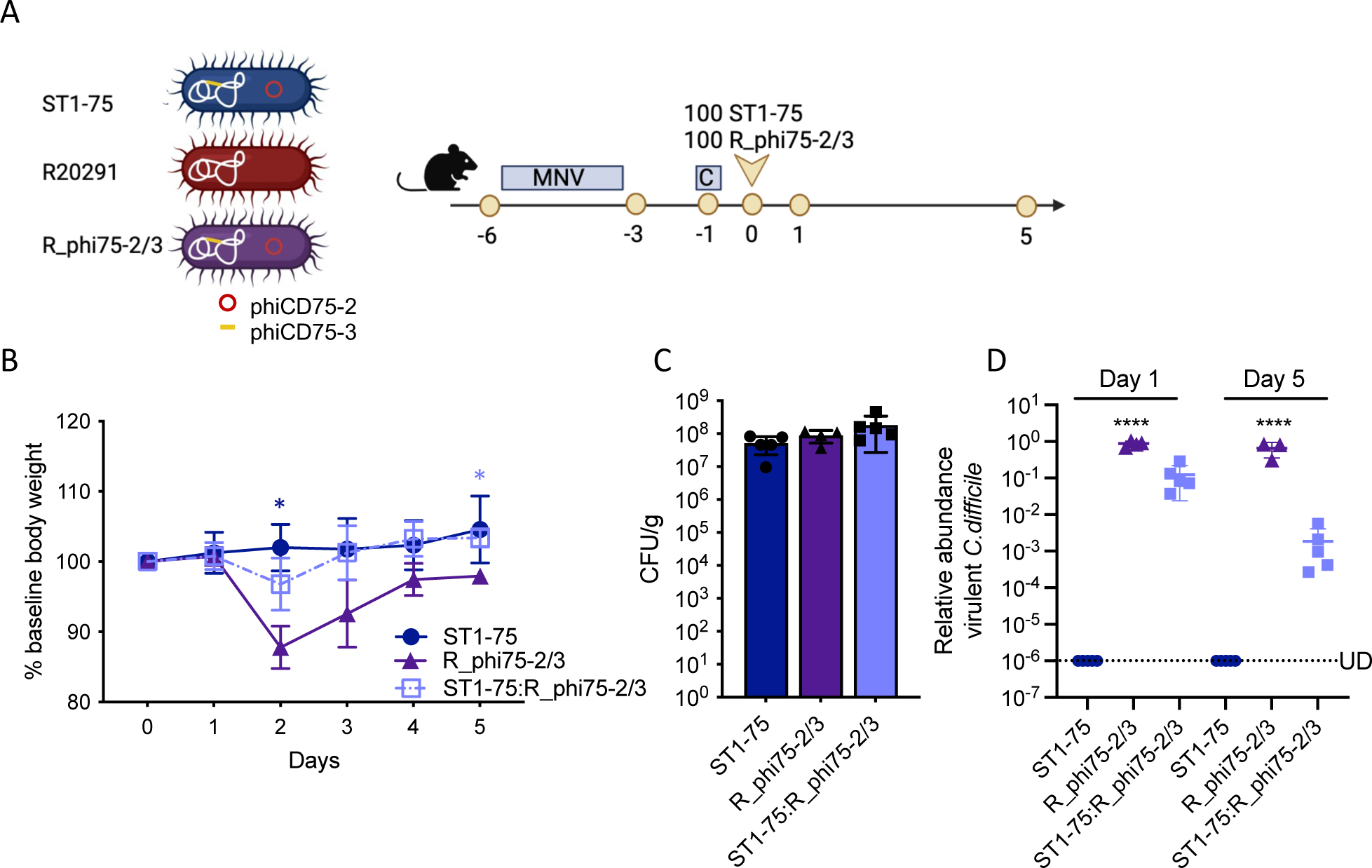
Protection by the avirulent ST1-75 isolate is not due to differences in lysogenic phages between strains. (A) Lysogeny R20291 strains generated to harbor phiCD75-2 and phiCD75-3 and infection procedure. Wild-type C57BL/6 mice (n = 4-5 per group) were treated with Metronidazole, Neomycin and Vancomycin (MNV, 0.25 g/L for each) in drinking water for 3 days, followed by one intraperitoneal injection of clindamycin (200 mg/mouse) 2 days after antibiotic recess. Then, mice were inoculated with 100 R_phi75-2/3 *C. difficile* spores and 100 ST1-75 *C. difficile* spores via oral gavage. Daily body weight was monitored for 5 days post-infection. (B) %Weight loss relative to the baseline of mice infected with indicated strains (C) Fecal colony-forming units measured by plating on selective agar on 1-day post-infection. (D) Relative abundance of R20291 in feces from infected mice was determined by measuring wildtype *cdtR* copies by qPCR on 1 day and 5 days post-infection. UD: Under the limit of detection. Statistical significance was calculated by unpaired t-test and One-way ANOVA, * p < 0.05, **** p <0.0001. Statistical significance was observed between R_phi75-2/3 to ST1-75 and ST1-75:R_phi75-2/3.

### Protection by the avirulent ST1-75 strain is not due to differences in the *cdtR* genes between strains

We previously showed that the avirulence of ST1-75 is due to a 69-base pair deletion in its *cdtR* gene, which encodes a regulator of CDT toxin, toxin A, and toxin B gene expression. Since introducing this *cdtR* mutation (*cdtR*\*) into R20291 is sufficient to render it avirulent (R20291 *cdtR*\* or R_ *cdtR*\*) (27), we sought to assess whether the *cdtR* mutation impacts the inter-strain competition between ST1-75 and R20291. To test this possibility, we coinfected mice with the avirulent R20291 *cdtR\** mutant we previously constructed (27) and the virulent R20291 parental strain (**Fig 3A**). Mice coinfected with R20291 *cdtR\** and R20291 lost weight to a similar degree as mice infected with R20291 alone, indicating that R20291 *cdtR\** is not as protective as ST1-75 (**Fig 3B**). When we measured the relative abundance of these two strains, we found that the R20291 *cdtR\** failed to outcompete R20291 by 1-day post-infection, in contrast with ST1-75. However, on 8 days post-infection, the R20291 *cdtR\** mutant outcompeted R20291, suggesting that the *cdtR\** mutation partially contributes to the competitive advantage of ST1-75 in *C. difficile* even though it cannot protect against virulent infection. Thus, ST1-75 must use additional mechanisms to outcompete R20291 more quickly and prevent mice from developing colitis.

**Figure 3.**
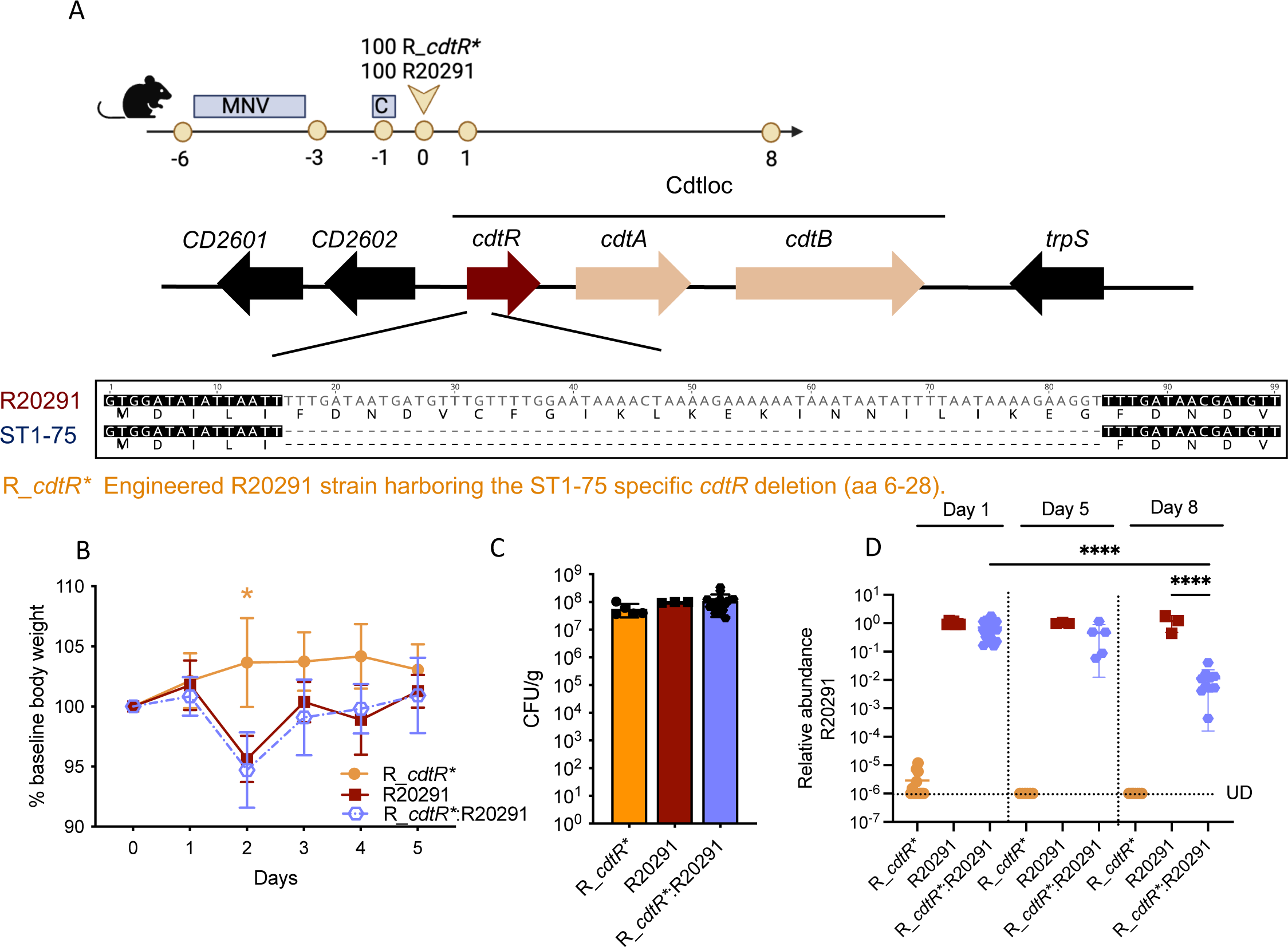
Coinfection with R20291 *cdtR\** does not protect against disease, although it outcompetes the parental R20291 strain. (A) Schematic of infection procedure and generating R20291 harboring 69-bp deletion in *cdtR.* Wild-type C57BL/6 mice (n = 3-5 per group) were treated with Metronidazole, Neomycin, and Vancomycin (MNV, 0.25 g/L for each) in drinking water for 3 days, followed by one intraperitoneal injection of clindamycin (200 mg/mouse) 2 days after antibiotic recess. Then, mice were inoculated with 100 R20291 *C. difficile* spores and 100 R_*cdtR* C. difficile* spores via oral gavage. Daily body weight was monitored for 5 days post-infection. (B) %Weight loss relative to the baseline of mice infected with indicated strains. (C) Fecal colony-forming units were measured by plating on selective agar on 1 day post-infection. (D) Relative abundance of R20291 in feces from infected mice was determined by measuring wildtype *cdtR* copies by qPCR on 1 day, 5 days, and 8 days post-infection. UD: Under the limit of detection. Statistical significance was calculated by unpaired t-test and One-way ANOVA, * p < 0.05, **** p <0.0001.

To gain further insight into this possibility, we compared the competitive fitness of R20291 *cdtR\** to the parental R20291 strain when R20291 *cdtR\** was used to infect mice 6 hours before R20291 infection. Despite being administered first, R20291 *cdtR*\* was unable to prevent R20291 from causing disease, even though the R20291 *cdtR\** strain eventually outcompeted its parental R20291 strain on Day 21 post-infection (**Fig S3B**). Since these clearance kinetics were slower than those observed for ST1-75, overall, the data suggest that ST1-75 has unique properties that allow it to outcompete R20291 and prevent virulent infection.

### Protection by the avirulent ST1-75 is independent of the microbiome and host immunity

To gain further insight into the mechanisms that contribute to ST1-75’s competitive advantage over R20291, we assessed whether the dysbiotic microbiome generated after MNV + Clindamycin treatment contributes to the competitive advantage of ST1-75 over R20291. Specifically, we measured the competitive fitness of ST1-75 and R20291 strains in germ-free mice during co-infection. Similar to the results obtained using antibiotic-treated mice, germ-free mice coinfected with ST1-75 and R20291 were protected from developing CDI (**Fig 4A-4B**), and the virulent R20291 strain was outcompeted by 24 hours post coinfection (**Fig 4C-4D**). These data indicate that the host microbiome is dispensable for ST1-75 to outcompete R20291 in mice. Additional experiments were conducted in MyD88^-/-^ mice, which lack a key innate immunity signaling factor (33), and Rag1^-/-^ mice, which lack adaptive immune responses (34). In both these immunodeficient strain backgrounds, the avirulent ST1-75 strains was able to outcompete the virulent R20291 strain (**data not shown**), indicating that the competitive advantage of ST1-75 in mice is independent of host immunity.

**Figure 4.**
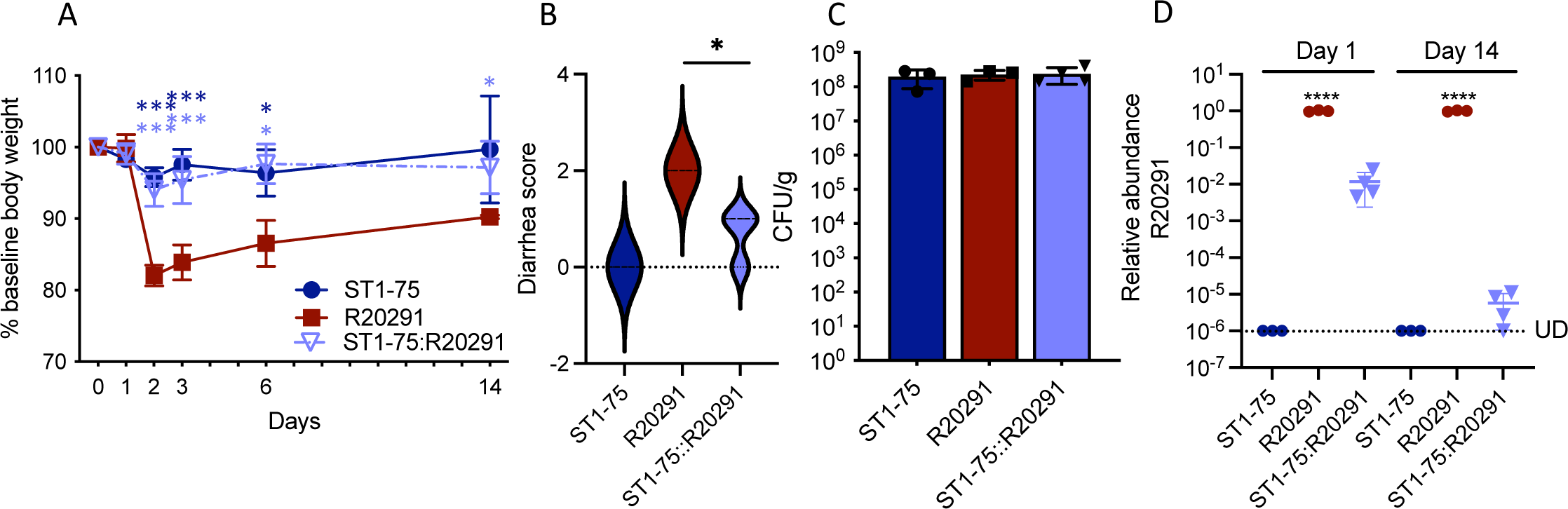
Protection by ST1-75 against virulent infection is independent of the microbiome. Germ-free mice (n=3-4 per group) were infected with ST1-75, R20291 or both at the same time at 200 total *C. difficile* spores. (A) %Weight loss relative to baseline of germ-free mice infected with indicated strains. (B) Diarrhea scores of mice infected with indicated strains 2 days post infection. (C) Fecal colony-forming units were measured by plating on selective agar on 1 day post-infection. (D) Relative abundance of R20291 in feces from infected mice was determined by measuring wildtype *cdtR* copies by qPCR on 1 day and 14 days post-infection. UD: Under the limit of detection. Statistical significance was calculated by unpaired t-test or One-way ANOVA, * p < 0.05, ** p < 0.01, **** p < 0.0001.

### Enhanced amino acid depletion provides ST1-75 competitive advantages to R20291

Since the ability of ST1-75 to outcompete R20291 in mice is independent of the murine microbiome or immune defenses, we compared the competitive fitness of these two strains in different growth conditions *in vitro*. First, we analyzed the relative fitness when ST1-75 and R20291 vegetative cells were co-inoculated at a 1:1 ratio in BHIS, a rich medium routinely used to culture *C. difficile*, at 3, 8, and 24 hrs post-co-inoculation. Surprisingly, ST1-75 did not gain a growth advantage over R20291 in rich broth, in contrast with our observations in mice (**Fig 5A**). When we co-cultured ST1-75 and R20291 in a medium derived from filtered cecal contents from germ-free mice, ST1-75 outcompeted R20291 by a ratio of 6:1 after 24 hrs and 12:1 after 96 hrs (**Fig 5B**). Conversely, when we compared the relative fitness of R20291 to the R20291 *cdtR\** in BHIS and *ex vivo* cecal content media, R20291 *cdtR\** exhibited similar fitness to R20291 in either BHIS or cecal media, consistent with the limited growth advantage of R20291 *cdtR*\* over R20291 observed in mice (**Fig 5A-5B**).

**Figure 5.**
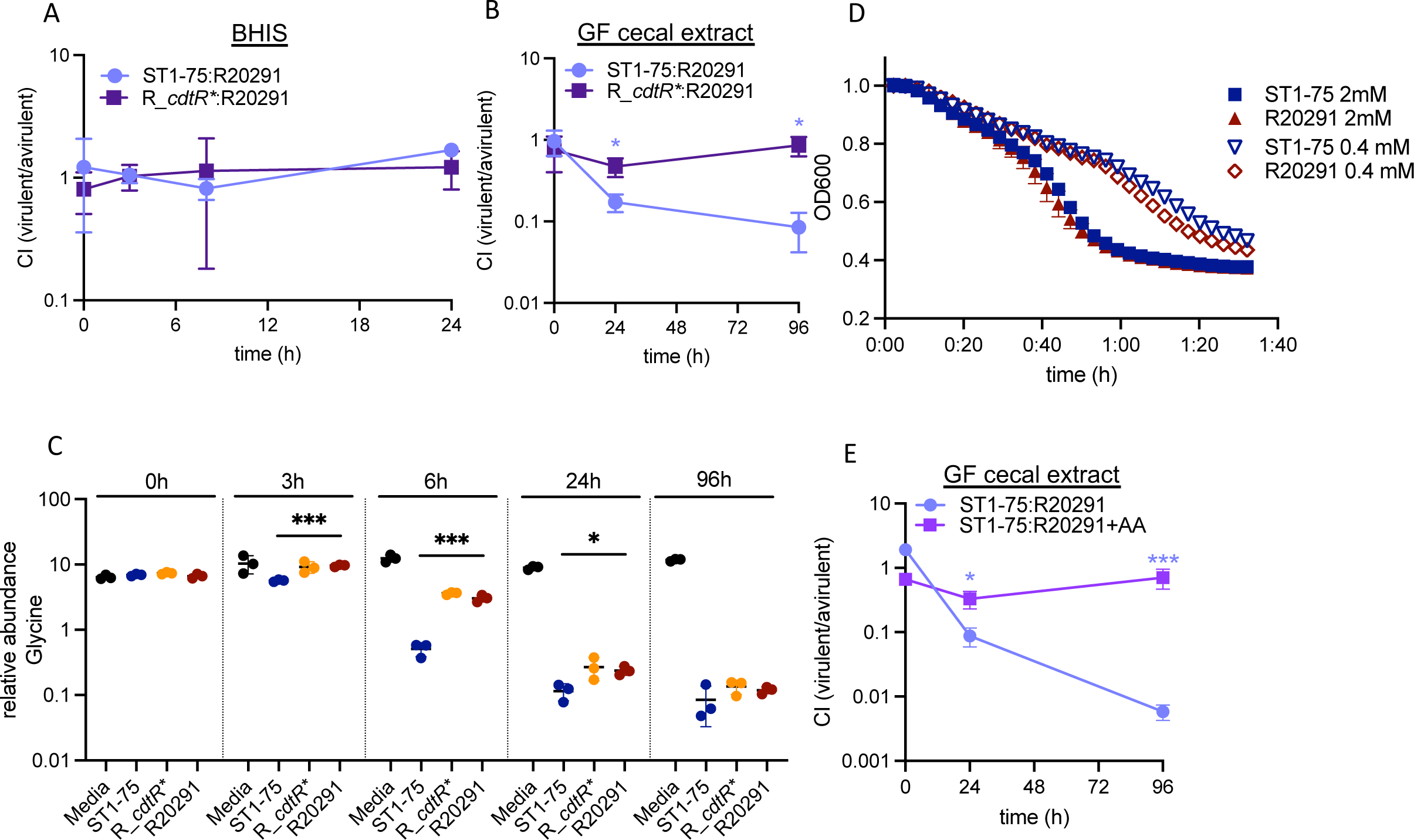
ST1-75 protection is attributable to faster amino acids depletion in nutrient-limited environment. (A-B, E) Relative abundance of R20291 in feces from infected mice was determined by measuring wildtype *cdtR* copies by qPCR. (A) Competitive index (CI) is shown by dividing the relative abundance of R20291 to avirulent strain (ST1-75/R_*cdtR*\*) in BHIS (B) Competitive index (CI) is shown by dividing the relative abundance of R20291 to avirulent strain (ST1-75/R_*cdtR*\*) in diluted germ-free cecal extract. (C) Metabolomic relative abundance of glycine in cecal extracts inoculated with either ST1-75, R_*cdtR\** or R20291 (D) OD-drop assay measuring germination defects of indicated strains while supplemented with 0.4 mM and 2 mM glycine. (E) Competitive index (CI) is shown by dividing the relative abundance of R20291 to avirulent strain ST1-75 in diluted germ-free cecal extract (spiked in with or without amino acids). Statistical significance was calculated by unpaired t-test or One-way ANOVA, * p < 0.5, *** p < 0.001, **** p < 0.0001.

Since cecal contents have relatively limited nutrients compared to rich BHIS medium, we hypothesized that the competition between ST1-75 and R20291 is nutrient-dependent. To test this hypothesis, we compared the contents of cecal cultures of ST1-75 vs. R20291 using metabolomics. These analyses revealed that several amino acids, including glycine, alanine, and phenylalanine, were depleted significantly faster by ST1-75 than either the WT or R20291 *cdtR\** strains (**Fig 5C and S4**).

Notably, glycine is a potent co-germinant for *C. difficile* spores (35), and previous work has shown that limiting amounts of glycine can impair the ability of a virulent *C. difficile* strain to colonize mice if the mice are pre-colonized with a low virulence *C. difficile* strain (23). This is because colonization by the low virulence strain depletes glycine from cecal contents and reduces the germination of the invading virulent *C. difficile* strain. Based on these observations, we considered the possibility that ST1-75 spores germinate more readily (i.e. are more sensitive to glycine co-germinant) than R20291 spores in the mouse gut such that ST1-75 can establish a replicative niche in the gut more rapidly than R20291 and thus reduce the availability of glycine and potentially R20291 spore germination. To test this possibility, we compared the sensitivity of ST1-75 and R20291 spores to low concentrations of glycine co-germinant. An optical density-based (OD_600_) germination assay was used to compare the co-germinant sensitivity of ST1-75 and R20291 at two physiologically relevant concentrations of glycine. These analyses revealed that the germination profiles for the ST1-75 and R20291 spores were identical at both glycine concentrations tested (**Fig 5D**), strongly suggesting that the competitive fitness advantage of ST1-75 is not due to differences in germination levels or rates.

On the other hand, several studies have implicated specific amino acids in promoting *C. difficile* colonization in the gut, consistent with the finding that CDI patients often have higher levels of amino acids than non-CDI patients (36, 37). To test if a limited availability of amino acids allows ST1-75 to outcompete R20291, we co-cultured ST1-75 and R20291 in cecal media supplemented with 18 amino acids at concentrations previously described to be sufficient for *C. difficile* growth (38). Amino acid supplementation was sufficient to ablate the growth advantage of ST1-75 over R20291 compared to the non-supplemented control (**Fig 5E**). These data strongly suggest that ST1-75 competes more efficiently for scarce amino acids in cecal contents and, by extension, during murine infection.

To identify additional amino acids beyond glycine, alanine, and phenylalanine (**Fig 5, S4**) that may impact the competition between ST1-75 and R20291, we measured the relative concentration of targeted amino acids after ST1-75 or R20291 were individually cultured in the amino acid-supplemented cecal media. These analyses revealed that valine, methionine, and isoleucine were also consumed faster by ST1-75 (**Fig S5**). Taken together, our data suggest that ST1-75 depletes multiple amino acids more rapidly than R20291, which presumably limits the growth of R20291 and allows ST1-75 to out-compete R20291 in the murine gut.

### ST1-75 confers long-term protection and can outcompete multiple virulent *C. difficile* strains

The avirulence of ST1-75 in both WT and immunodeficient mice (27) and its ability to outcompete a virulent *C. difficile* strain suggest that it could be used as a bacterial therapeutic for preventing CDI. However, a few concerns are routinely raised when considering the use of probiotics for providing colonization resistance against pathogens, specifically how long probiotics colonize the host gut and how well they persist upon antibiotic treatment. To address these concerns, we inoculated antibiotic-treated mice with ST1-75 (one oral gavage) and monitored its long-term colonization (one month) (**Fig 6A**). We found that ST1-75 established high-level colonization throughout the month (**Fig 6B**). After this colonization period, we treated ST1-75 colonized mice with additional antibiotics and then challenged them with virulent R20291 (**Fig 6A**). While the antibiotics successfully reduced ST1-75 levels by ∼3-log (**Fig 6B**), the CFUs of ST1-75 rebounded prior to the R20291 challenge. This rebound was sufficient to provide colonization resistance against R20291 infection and protect mice from CDI colitis (**Fig 6C-6E**). Such “relapse” of ST1-75 is likely attributable to its antibiotic-resistant spores. These data are promising and support the potential of using ST1-75 to confer long-term colonization resistance and protection against *C. difficile* in patients following repeat antibiotic treatment. Mouse experiments are not fully translatable to humans as mice are coprophagic, allowing them to re-inoculate ST1-75 on a regular basis. Thus, in a non-coprophagic recipient, serial oral administration may be required to maintain colonization resistance.

**Figure 6.**
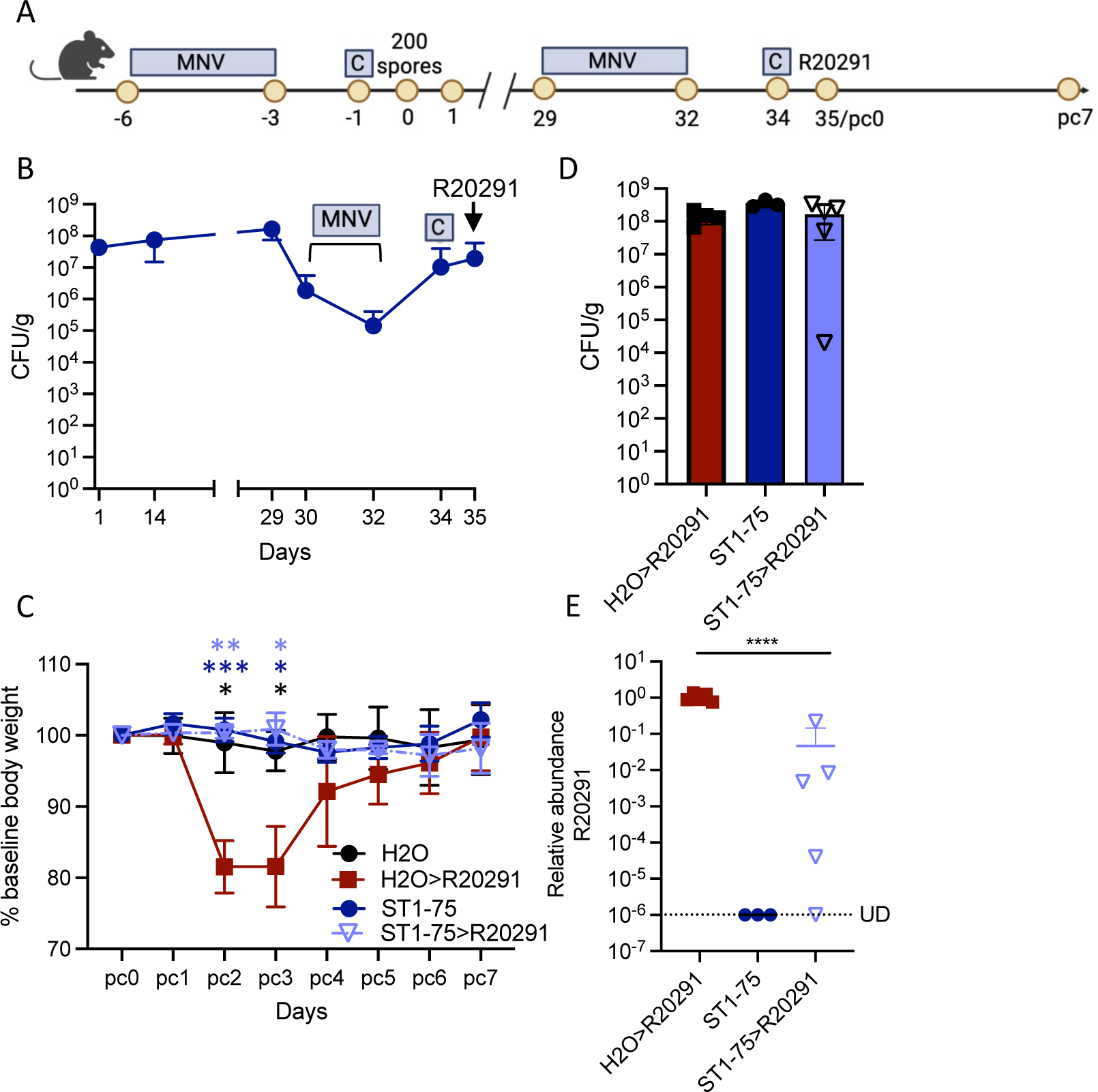
ST1-75 confers long-term protection via colonization resistance. (A) Schematic of the experimental procedure. Wild-type C57BL/6 mice (n = 3-5 per group) were treated with Metronidazole, Neomycin and Vancomycin (MNV, 0.25 g/L for each) in drinking water for 3 days, followed by one intraperitoneal injection of clindamycin (200 mg/mouse) 2 days after antibiotic recess. Then, mice were inoculated with 200 ST1-75 *C. difficile* spores via oral gavage. Infected mice were maintained for a month and then treated with MNV and Clindamycin again before challenging with 200 R20291 *C. difficile* spores. Daily body weight was monitored for 7 days post R20291 challenge. (B) Fecal colony-forming units were measured by plating on selective agar on indicated days post-infection. (C) %Weight loss relative to the baseline of mice infected with indicated strains. (D) Fecal colony-forming units were measured by plating on selective agar on 1 day post-R20291 infection. (E) Relative abundance of R20291 in feces from infected mice was determined by measuring wildtype *cdtR* copies by qPCR on 1 day post-R20291 infection. UD: Under the limit of detection. Statistical significance was calculated by unpaired t-test or One-way ANOVA, * p < 0.05, ** p < 0.01, **** p < 0.0001.

We next wondered whether ST1-75 can outcompete *C. difficile* strains that were isolated more recently or are from other ribotypes. Using a similar coinfection strategy by coinfecting antibiotic-treated mice with a 1:1 ratio of ST1-75 and either ST1-12 or ST1-49 (**Fig 7A**). ST1-12 and ST1-49 are recent isolates that we previously reported to be more virulent than R20291 in mice (27). Coinfection with ST1-75 reduced the disease severity caused by ST1-49 and, to a lesser extent, ST1-12 (**Fig 7B**), although the data did not reach statistical significance after adjusting the false discovery rate for multiple comparisons. Indeed, co-inoculating ST1-75 and ST1-49 almost completely reversed the weight loss phenotype (**Fig 7B**). qPCR measuring the relative abundance of each *C. difficile* strain indicated that ST1-75 rapidly outcompetes ST1-49, consistent with the weight loss phenotype (**Fig 7C-7D**). In contrast, ST1-75 was unable to clear ST1-12 by day 1, consistent with the weight loss observed in mice coinfected with ST1-75 and ST1-12 (**Fig 7B-7D**). However, ST1-12 was eventually outcompeted by ST1-75 at later time points (**Fig 7D**).

**Figure 7.**
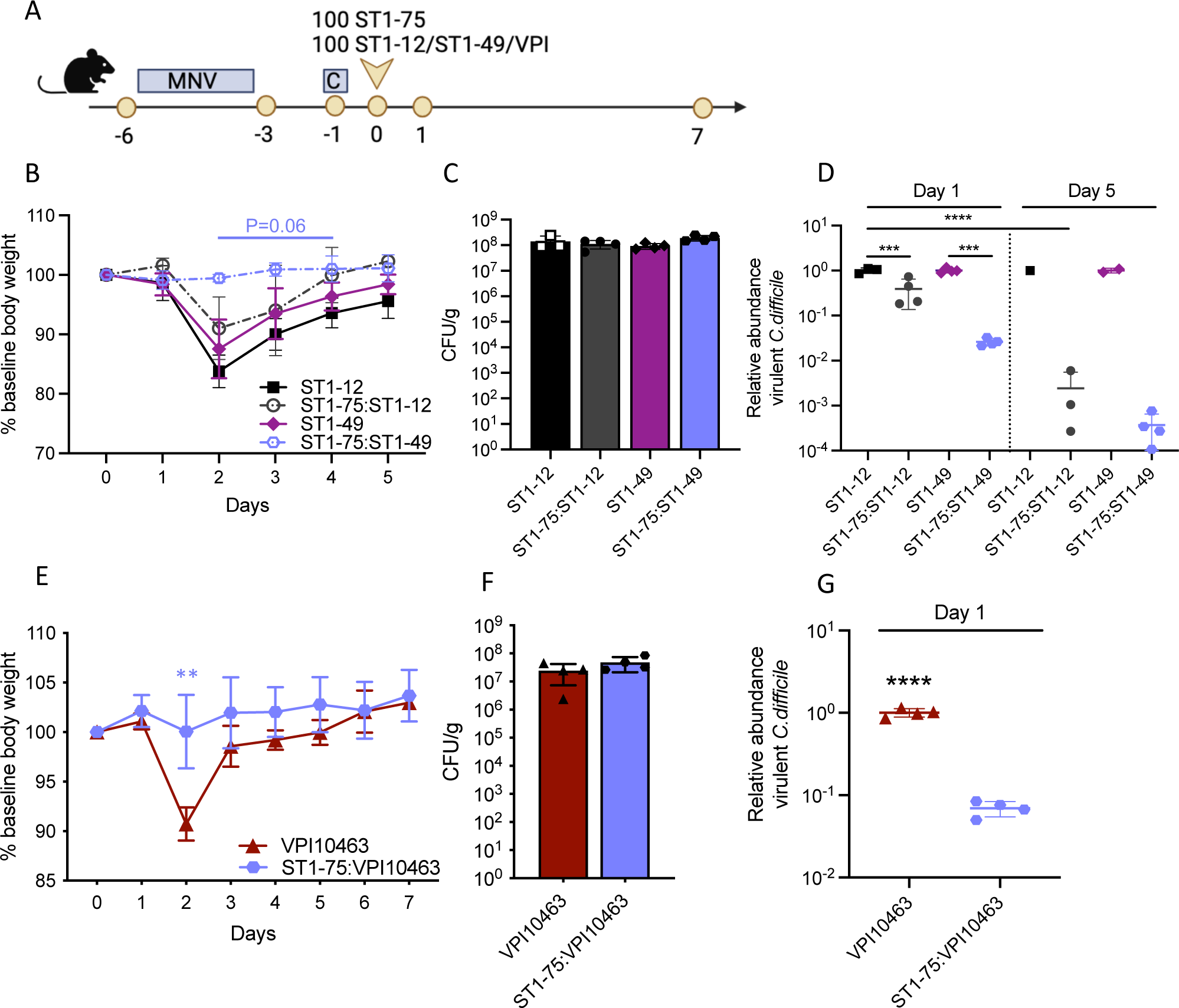
ST1-75 clears recent clinical ST1 isolates and the distantly related strain VPI10463. (A) Schematic of the experimental procedure. Wild-type C57BL/6 mice (n = 4 per group) were treated with Metronidazole, Neomycin and Vancomycin (MNV, 0.25 g/L for each) in drinking water for 3 days, followed by one intraperitoneal injection of clindamycin (200 mg/mouse) 2 days after antibiotic recess. Then, mice were inoculated with 100 ST1-75 *C. difficile* spores and 100 either ST1-12, ST1-49 or VPI10463 *C. difficile* spores via oral gavage. Daily body weight was monitored for 5-7 days post-infection. (B, E) %Weight loss relative to the baseline of mice infected with indicated strains. (C, F) Fecal colony-forming units were measured by plating on selective agar on 1 day post-infection. (D, G) Wildtype *cdtR* copies in feces from infected mice were measured by qPCR on 1 day and 5 days post infection. Some mice dead or did not produce a fecal pellet on day 5. Statistical significance was calculated by unpaired t-test or One-way ANOVA, ** p < 0.01, *** p < 0.001, **** p < 0.0001.

VPI10463 strain is commonly used to evaluate therapeutics since it is a high toxin producer and cause lethal infections in mice (39, 40). To determine whether ST1-75 could protect against non-RT027 ribotype infection, we coinfected antibiotic-treated mice with 1:1 ratio of ST1-75 and VPI10463. Notably, ST1-75 also outcompetes this more distantly related *C. difficile* strain (41) and protects mice from its virulence (**Fig 7E-7G**). Overall, these data suggest that ST1-75 has a competitive advantage over closely and more distantly related *C. difficile* strains.

## Discussion

While FMT is effective at treating many recurrent *C. difficile* infections, the high cost and challenges in ensuring reproducibility and production at scale has motivated the search for more defined bacterial therapeutics. Several studies have shown that avirulent or less virulent *C. difficile* strains can prevent lethal CDI, yet we do not know all the mechanisms underlying the protection (23, 25, 42, 43). Herein, we characterized a toxigenic but avirulent *C. difficile* isolate, ST1-75, that protects against virulent CDI by outcompeting virulent *C. difficile* strains. ST1-75 can outcompete several closely or distantly related virulent strains, including R20291 (RT027), ST1-12 (RT027), ST1-49 (RT027), and VPI10463 (RT087). Our analyses revealed that the ability of ST1-75 to outcompete these more virulent strains can be attributed to ST1-75’s enhanced capacity to consume amino acids, which limits the growth of the virulent strains. Finally, we show that ST1-75 establishes long-term colonization in the host and persists after subsequent antibiotic treatment, suggesting a potential prophylactic capacity to prevent primary and recurrent CDI in patients with antibiotic-induced dysbiosis.

ST1-75 outcompeted virulent R20291 when administered at a 1:1 ratio during co-infection. ST1-75’s efficacy at low doses is significant because previous non-toxigenic and low-virulence *C. difficile* strains that have been shown to confer protection against virulent *C. difficile* infections in animals required that they be administered at a significantly higher dosage over virulent *C. difficile* strains and/or days before the virulent strain challenge (23, 42). For example, while pre-inoculating nontoxigenic *C. difficile* demonstrated protection, coinfecting nontoxigenic *C. difficile* could not compete with virulent strains, leading to the mortality of infected hamsters (44). In another study, a low-virulence *C. difficile* strain CD630 required 100-fold higher inoculum to protect mice from virulent VPI10463 challenge (23). These studies suggest that such nontoxigenic or low-virulence strains may have reduced competitive fitness over virulent *C. difficile* strains.

The competitive advantages of ST1-75 over the other nontoxigenic or low-virulence *C. difficile* strains may be attributable to its mechanisms to outcompete virulent strains. While the mechanisms utilized by non- or low-virulence strains to provide colonization resistance against more virulent *C. difficile* strains still need further exploration, a previous study reported that pre-colonizing mice with low-virulence *C. difficile* depletes the essential germinant glycine to restrict the germination of the secondary virulent strain (23). Here, we found that ST1-75 competes with virulent strains in a different manner, specifically by quickly consuming available amino acids to limit the growth of the virulent strains. Our metabolomic analyses indicate a few amino acid candidates that may be important for ST1-75 to outcompete R20291, namely alanine, phenylalanine, valine, methionine, and isoleucine, that are depleted faster by ST1-75 than R20291. These include important amino acids involved in Stickland metabolism, which were previously implicated to play important roles in *C. difficile* colonization in the gut (40). Here, by supplementing amino acids in a scarce medium, we provided direct evidence supporting the importance of amino acid availability in impacting inter-stain competition. Ornithine and proline metabolisms were previously shown as a battle ground between commensal microbiome and *C. difficile*, with *C. difficile* unable to metabolize ornithine or proline colonizing poorly (36, 45). Our assay did not detect ornithine and proline was not differentially depleted by ST1-75 vs. R20291. Future experiments can dissect if a specific amino acid or a group of amino acids would be needed during the strain-strain competition between ST1-75 and R20291. Such information can contribute to the design of adjunctive therapies, including diet modifications. Additional investigation should also focus on determining the molecular basis of ST1-75’s enhanced capacity to consume such amino acids.

During the investigation of the dynamics of ST1-75 and R20291 competition during coinfection, we found that administering ST1-75 6 hrs post R20291 infection at high dose (50-fold over R20291) outcompeted R20291 by 24 hours, similar to the simultaneous coinfection. However, ST1-75 administration 6 hrs after R20291 did not reverse the level of weight loss. We speculate that there are relatively early changes in intestinal environment, caused by R20291, that commit to pathogenesis. Depending on types of antibiotics treatment, *C. difficile* CFU in the colon and feces can be detected around 6-12 hrs post-infection while spores and toxins were detected 18-24 hrs post-infection (46). The fulminant weight loss can only be observed 2-3 days post-infection in most reported mouse experiments (34, 39, 47). Due to this “delayed” phenotype, most studies on immune changes focused on these time points (34, 48) while the earlier changes of the *C. difficile* infected gut were not well-investigated. We hypothesize that the early events in the gut during *C. difficile* infection contribute to the pathogenesis trajectory.

Our finding that the R20291 *cdtR\** mutant has a small competitive fitness over R20291 is interesting. CdtR is the response regulator of *C. difficile* binary toxin CDT, and mutations in the *cdtR* gene lead to reduced production of CDT and primary toxins TcdA and TcdB (27). The roles of toxins during *C. difficile* strain-strain competition remains unresolved and additional studies are required to determine whether the fitness of R20291 *cdtR\** mutant is attributable to its reduced toxin production. Previous studies, however, showed that *C. difficile* producing no toxin (NTCD) have reduced fitness compared to virulent strains, while *C. difficile* producing some toxins (ST1-75, LEM1) have greater fitness (24). We have yet to determine whether toxins enhance the growth advantage of ST1-75, especially since toxins were reported to liberate nutrients from the mucus and enhance *C. difficile* colonization (36, 49). Future efforts should also be made to determine if toxin production (though at a reduced level due to the *cdtR* mutation) of ST1-75 is dispensable for its competitive fitness. This is important to know before one can further optimize ST1-75 by knocking out all its toxin genes to make a safer biotherapeutic candidate.

Strategies to use a non-virulent, commensal bacterial strain to compete for necessary nutrients and thus protect against virulent infections were also demonstrated for a couple of other bacterial pathogens. For example, a *Klebsiella sp.* was first shown to enhance resistance to *Enterobacteriaceae*, such as *E. coli* and *Salmonella* via galactitol competition (50). Galactitol competition was also observed while using a commensal *E. coli* strain Mt1B1 to block *Salmonella* Typhimurium invasion (51). Another study demonstrated that *Klebsiella oxytoca* outcompetes *Klebsiella pneumoniae* for various aromatic and non-aromatic beta-glucosides (52). The advantages of using an avirulent strain of a bacterial pathogen to prevent virulent infection are that they would occupy the same niche and compete with the most overlapping nutritional repertoire. Further investigations are needed to better understand the inter-strain competitions to improve the safety and efficacy of using avirulent “pathogens” clinically.

In summary, we described the use of avirulent isolate ST1-75 as a prophylactic to prevent virulent CDI in mice. ST1-75 can outcompete virulent strains at a low dosage (1:1 ratio) and during coinfection, granting it high potential in preventing CDI recurrence and transmission. The mechanism we have discovered is that ST1-75 depletes specific amino acids more quickly than R20291, thus limiting such nutritional access by the virulent strain. Overall, our study provides a unique mechanism for *C. difficile* strain-strain competition that may be applied to other bacterial-bacterial interactions and support the use of ST1-75 as a potential biotherapeutic in preventing colonization and disease caused by virulent *C. difficile*.

## Material and Methods

### Mouse husbandry

Wild-type C57BL/6J mice, aged between 6 to 8 weeks, were purchased from the Jackson Laboratories. Germ-free C57Bl/6J mice were bred and maintained in plastic gnotobiotic isolators within the University of Chicago Gnotobiotic Core Facility and fed ad libitum autoclaved standard chow diet (LabDiets 5K67) before transferring to BSL2 room for infection. Mice housed in the BSL2 animal room are fed irradiated feed (Envigo 2918) and provided with acidified water. All mouse experiments were performed in compliance with University of Chicago’s institutional guidelines and were approved by its Institutional Animal Care and Use Committee.

### *C. difficile* spore preparation and numeration

*C. difficile* sporulation and preparation was processed as described previously (27). Briefly, single colonies of *C. difficile* strains were inoculated in deoxygenated BHIS broth and incubated anaerobically for 40-50 days. *C. difficile* cells were harvested by centrifugation and five washes with ice-cold water. The cells were then suspended in 20% (w/v) HistoDenz (Sigma, St. Louis, MO) and layered onto a 50% (w/v) HistoDenz solution before centrifugating at 15,000 × g for 15 minutes to separate spores from vegetative cells. The purified spores pelleted at the bottom were then collected and washed for four times with ice-cold water to remove traces of HistoDenz, and finally resuspended in sterile water. Prepared spores were heated to 60°C for 20 min to kill vegetative cells, diluted and plated on both BHIS agar and BHIS agar containing 0.1% (w/v) taurocholic acid (BHIS-TA) for numeration. Spore stocks for mouse infection were verified to have less than 1 vegetative cell per 200 spores (as the infection dose).

For OD_600_ kinetics assays, spores were prepared a little differently (53). Briefly, single colonies were used to inoculate liquid BHIS cultures, which were grown to early stationary phase before being back diluted 1:50 into BHIS. When the cultures reached an OD_600_ between 0.35 and 0.75, 120 μL of this culture were spread onto 70:30 agar plates and sporulation was induced as previously described for 5 days. The spores were then harvested into ice-cold, sterile water, washed 6 times in ice-cold water and incubated overnight in water at 4°C. The following day, the samples were pelleted and treated with DNase I (New England Biolabs) at 37°C for 60 minutes and purified on a 20%/50% HistoDenz (Sigma Aldrich) gradient. The resulting spores were washed again 2–3 times in water, and spore purity was assessed using phase-contrast microscopy (>95% pure). The optical density of the spore stock was measured at OD_600_, and spores were stored in water at 4°C.

### Virulence assessment of clinical isolates in mice

Mice were treated and infected with *C. difficile* spores as described previously (27). Briefly, SPF mice were treated with antibiotic cocktail containing metronidazole, neomycin and vancomycin (MNV) in drinking water (0.25g/L for each antibiotic) for 3 days, 2 days after removing MNV, the mice received one dose of clindamycin (200 µg/mouse) via intraperitoneal injection. Mice were then infected the next day with 200 *C. difficile* spores via oral gavage. Germ-free mice were infected with 200 *C. difficile* spores via oral gavage without antibiotic treatments. Following infection, mice were monitored and scored for disease severity by four parameters (34): weight loss (> 95% of initial weight = 0, 95%–90% initial weight = 1, 90%–80% initial weight = 2, < 80% = 3), surface body temperature (> 95% of initial temp= 0, 95%–90% initial temp = 1, 90%–85% initial temp = 2, < 85% = 3), diarrhea severity (formed pellets = 0, loose pellets = 1, liquid discharge = 2, no pellets/caked to fur = 3), morbidity (score of 1 for each symptoms with max score of 3; ruffled fur, hunched back, lethargy, ocular discharge).

### Quantification of fecal colony forming units

Fecal pellets from *C. difficile* infected mice were harvested and resuspended in deoxygenated phosphate-buffed saline (PBS), diluted and plated on BHI agar supplemented with yeast extract, taurocholic acid, L-cysteine, D-cycloserine and cefoxitin (CC-BHIS-TA) at 37°C anaerobically for overnight (27, 48).

### Preparation of Germ-Free Cecal Extract Media

Cecal contents from germ-free wild type C57BL/6 mice were harvested and resuspended in sterile water (200 mg cecal content per 1 mL water). The cecal slurry was vortexed and centrifuged at 4,300 x g using an Allegra X-14R Centrifuge (Beckman Coulter) for 10 minutes to pellet solid waste material. The supernatant was collected and centrifuged 3 more times before filtration through a 0.2-micron filter. The filter-sterilized cecal supernatant media was aliquoted and frozen at −20°C.

### Amino Acid Supplementation

Fresh amino acid stocks (histidine, glycine, arginine, phenylalanine, methionine, threonine, alanine, lysine, serine, valine, isoleucine, leucine, proline, aspartic acid, glutamic acid, tyrosine, cysteine) were prepared by suspending 10 mg of each compound in 10 mL of 5N KOH and freezing 1 mL aliquots at −20°C. Amino acids were added to thawed germ-free cecal supernatant media at concentrations previously described to be sufficient for *C. difficile* growth (38). Otherwise, an equal volume of sterile water was added to the cecal media. Then, each condition was pH adjusted to 8.4-8.5, filtered through a 0.2-micron filter, and reduced in an anaerobic chamber overnight.

### C. difficile culturing

In an anaerobic chamber (Coylabs), *C. difficile* strains were streaked out on BHI agar supplemented with yeast extract (BHIS) containing 0.1% (w/v) taurocholic acid (BHIS-TA) for single colonies. Colonies were then picked for overnight growth in BHIS broth at 37°C anaerobically. Overnight cultures were 1:10 diluted in BHIS and incubated for 3-4 hours to grow to the exponential phase, then normalized to OD 0.05 as the starting point for time course incubation, in either BHIS or cecal extracts. All media and cecal extract were deoxygenated in anaerobic chamber before use.

### Metabolite Extraction from Plasma/Serum/Culture Supernatant

Samples were incubated at −80°C for at least one hour, or up to overnight. Extraction solvent (4 volumes of 100% methanol spiked with internal standards and stored at −80°C) was added to the liquid sample (1 volume) in a microcentrifuge tube. Tubes were then centrifuged at −10°C, 20,000 x g for 15 min and supernatant was used for subsequent metabolomic analysis.

### Metabolite Analysis using GC-nCI-MS and PFBBr Derivatization

Short chain fatty acids were derivatized as described by Haak et al (54). with the following modifications. The metabolite extract (100 μL) was added to 100 μL of 100 mM borate buffer (pH=10), (Thermo Fisher, 28341), 400 μL of 100 mM pentafluorobenzyl bromide, (Millipore Sigma; 90257) in acetonitrile, (Fisher; A955-4), and 400 μL of n-hexane (Acros Organics; 160780010) in a capped mass spec autosampler vial (Microliter; 09-1200). Samples were heated in a thermomixer C (Eppendorf) to 65°C for 1 hour while shaking at 1300 rpm. After cooling to room temperature, samples were centrifuged at 4°C, 2000 x g for 5 min, allowing phase separation. The hexanes phase (100 μL) (top layer) was transferred to an autosampler vial containing a glass insert and the vial was sealed. Another 100 μL of the hexanes phase was diluted with 900 μL of n-hexane in an autosampler vial. Concentrated and dilute samples were analyzed using a GC-MS (Agilent 7890A GC system, Agilent 5975C MS detector) operating in negative chemical ionization mode, using a HP-5MSUI column (30 m x 0.25 mm, 0.25 μm; Agilent Technologies 19091S-433UI), methane as the reagent gas (99.999% pure) and 1 μL split injection (1:10 split ratio). Oven ramp parameters: 1 min hold at 60°C, 25°C per min up to 300°C with a 2.5 min hold at 300°C. Inlet temperature was 280°C and transfer line was 310°C. A 10-point calibration curve was prepared with acetate (100 mM), propionate (25 mM), butyrate (12.5 mM), and succinate (50 mM), with 9 subsequent 2x serial dilutions. Data analysis was performed using MassHunter Quantitative Analysis software (version B.10, Agilent Technologies) and confirmed by comparison to authentic standards. Normalized peak areas were calculated by dividing raw peak areas of targeted analytes by averaged raw peak areas of internal standards. Raw data (.d, Agilent) was converted with mzMine to open-source formatting (.mzML) and uploaded to MassIVE.

### OD_600_ kinetics assays

OD_600_ kinetics assays were conducted as described previously with minor modifications (53). For OD_600_ kinetics assays with varying concentrations of glycine, ∼2.3 x 10^7^ spores (0.8 OD_600_ units) for each condition tested were resuspended in either 1X 50 mM HEPES, 100 mM NaCl, pH=8 and aliquoted into a well of a 96-well plate for each condition tested. 5-fold serial dilutions of 100 mM glycine were added to spores resuspended in the appropriate buffer. The spores were then exposed to 1% taurocholate (19 mM) in a total volume of 200 μL and the OD_600_ of the samples was measured every 3 minutes in a Synergy H1 microplate reader (Biotek) at 37°C with constant shaking between readings. The change in OD_600_ over time was calculated as the ratio of the OD_600_ at each time point to the OD_600_ at time zero. All assays were performed on three independent spore preparations.

### DNA extraction and quantitative polymerase chain reaction (qPCR)

Fecal DNA was extracted using DNeasy PowerSoil Pro Kit or QIAamp PowerFecal Pro kit (Qiagen) according to the manufacturer’s instructions. Quantitative PCR was performed on genomic DNA using primers:

cdtR_commonFor 5’-TCTCCTAGTGTTATTTTACGTATTTTT-3’

cdtR_wtFor 5’-TGTTTGTTTTGGAATAAAACTAAAAGA-3’

VPI_cdtR_commonFor 5’-TCTCCTAGTGTTATTTTATGTATTTT-3’

VPI_cdtR_wtFor 5’-TGTTTGTTTTGGAATGAAACTAAAAGA-3’

cdtR_Rev 5’-TTGTGCTATCCATAATCCATCACA-3’

with PowerTrack SYBR Green Master Mix (Thermo Fisher). Reactions were run on a QuantStudio 6 pro (Thermo Fisher). Relative abundance was normalized by ΔΔCt.

### Quantification and statistical analysis

Results represent means ± SD. Statistical significance was determined by t test and one-way ANOVA test. Multiple comparisons were corrected with False Discovery Rate with desired FDR at 0.05. Statistical analyses were performed using Prism GraphPad software v9.3.1 (∗ p < 0.05, ∗∗ p < 0.01, ∗∗∗ p < 0.001, ∗∗∗∗ p < 0.0001)

## Acknowledgements

We thank Kristin Kolar, Emma Elmiger and the animal facilities of the University of Chicago for their help on mouse experiments. We thank the Pamer lab members and Shen lab members for helpful discussions. This work was supported by the National Institutes of Health R01 AI095706 and by the Duchossois Family Institute of the University of Chicago. The funders had no role in study design, data collection and interpretation, or the decision to submit the work for publication. Schematics were created with bioRender.com.

## Author contributions

Q.D., L.C.F. A.S. and E.G.P conceived the project. Q.D., S.H., E.M., S.S.S., and H.L. analyzed the data. Q.D., S.H., E.M., R.C.S., M-M.A., C.M., C.W., V.B., A.S., A.R., M.M., D.M., J.L., M.M., and A.S. performed experiments. Q.D., A.S., L.C.F., and E.G.P interpreted the results and wrote the manuscript.

## References

1 . Cdc A. 2019. Antibiotic resistance threats in the United States. US Dep Health Hum Serv Wash DC USA.

2. Abt MC, McKenney PT, Pamer EG. 2016. *Clostridium difficil*e colitis: pathogenesis and host defence. Nat Rev Microbiol 14:609–620.

3. Sorbara MT, Pamer EG. 2019. Interbacterial mechanisms of colonization resistance and the strategies pathogens use to overcome them. Mucosal Immunol 12:1–9.

4. Feuerstadt P, Boules M, Stong L, Dahdal DN, Sacks NC, Lang K, Nelson WW. 2021. Clinical complications in patients with primary and recurrent *Clostridioides difficile* infection: A real-world data analysis. SAGE Open Med 9:2050312120986733.

5. Turner NA, Anderson DJ. 2020. Hospital Infection Control: *Clostridioides difficile*. Clin Colon Rectal Surg 33:98–108.

6. Pokrywka M, Buraczewski M, Frank D, Dixon H, Ferrelli J, Shutt K, Yassin M. 2017. Can improving patient hand hygiene impact *Clostridium difficile* infection events at an academic medical center? Am J Infect Control 45:959–963.

7. Barker AK, Zellmer C, Tischendorf J, Duster M, Valentine S, Wright MO, Safdar N. 2017. On the hands of patients with *Clostridium difficile*: A study of spore prevalence and the effect of hand hygiene on C difficile removal. Am J Infect Control 45:1154–1156.

8. Kaier K, Hagist C, Frank U, Conrad A, Meyer E. 2009. Two time-series analyses of the impact of antibiotic consumption and alcohol-based hand disinfection on the incidences of nosocomial methicillin-resistant Staphylococcus aureus infection and *Clostridium difficile* infection. Infect Control Hosp Epidemiol 30:346–353.

9. Vernaz N, Sax H, Pittet D, Bonnabry P, Schrenzel J, Harbarth S. 2008. Temporal effects of antibiotic use and hand rub consumption on the incidence of MRSA and *Clostridium difficile*. J Antimicrob Chemother 62:601–607.

10. Boyce JM, Guercia KA, Sullivan L, Havill NL, Fekieta R, Kozakiewicz J, Goffman D. 2017. Prospective cluster controlled crossover trial to compare the impact of an improved hydrogen peroxide disinfectant and a quaternary ammonium-based disinfectant on surface contamination and health care outcomes. Am J Infect Control 45:1006–1010.

11. Wenisch C, Parschalk B, Hasenhündl M, Hirschl AM, Graninger W. 1996. Comparison of vancomycin, teicoplanin, metronidazole, and fusidic acid for the treatment of *Clostridium difficile*-associated diarrhea. Clin Infect Dis Off Publ Infect Dis Soc Am 22:813–818.

12. DiDiodato G, McArthur L. 2016. Evaluating the Effectiveness of an Antimicrobial Stewardship Program on Reducing the Incidence Rate of Healthcare-Associated *Clostridium difficile* Infection: A Non-Randomized, Stepped Wedge, Single-Site, Observational Study. PloS One 11:e0157671.

13. Patton A, Davey P, Harbarth S, Nathwani D, Sneddon J, Marwick CA. 2018. Impact of antimicrobial stewardship interventions on *Clostridium difficile* infection and clinical outcomes: segmented regression analyses. J Antimicrob Chemother 73:517– 526.

14. Baur D, Gladstone BP, Burkert F, Carrara E, Foschi F, Döbele S, Tacconelli E. 2017. Effect of antibiotic stewardship on the incidence of infection and colonisation with antibiotic-resistant bacteria and *Clostridium difficile* infection: a systematic review and meta-analysis. Lancet Infect Dis 17:990–1001.

15. Feazel LM, Malhotra A, Perencevich EN, Kaboli P, Diekema DJ, Schweizer ML. 2014. Effect of antibiotic stewardship programmes on *Clostridium difficile* incidence: a systematic review and meta-analysis. J Antimicrob Chemother 69:1748–1754.

16. de Bruyn G, Gordon DL, Steiner T, Tambyah P, Cosgrove C, Martens M, Bassily E, Chan E-S, Patel D, Chen J, Torre-Cisneros J, Fernando De Magalhães Francesconi C, Gesser R, Jeanfreau R, Launay O, Laot T, Morfin-Otero R, Oviedo-Orta E, Park YS, Piazza FM, Rehm C, Rivas E, Self S, Gurunathan S. 2021. Safety, immunogenicity, and efficacy of a *Clostridioides difficile* toxoid vaccine candidate: a phase 3 multicentre, observer-blind, randomised, controlled trial. Lancet Infect Dis 21:252–262.

17. Pfizer. 2023. A PHASE 3, PLACEBO-CONTROLLED, RANDOMIZED, OBSERVER-BLINDED STUDY TO EVALUATE THE EFFICACY, SAFETY, AND TOLERABILITY OF A *CLOSTRIDIUM DIFFICILE* VACCINE IN ADULTS 50 YEARS OF AGE AND OLDER. NCT03090191. Clinical trial registration. clinicaltrials.gov.

18. Khanna S, Assi M, Lee C, Yoho D, Louie T, Knapple W, Aguilar H, Garcia-Diaz J, Wang GP, Berry SM, Marion J, Su X, Braun T, Bancke L, Feuerstadt P. 2022. Efficacy and Safety of RBX2660 in PUNCH CD3, a Phase III, Randomized, Double-Blind, Placebo-Controlled Trial with a Bayesian Primary Analysis for the Prevention of Recurrent *Clostridioides difficile* Infection. Drugs 82:1527–1538.

19. Feuerstadt P, Louie TJ, Lashner B, Wang EEL, Diao L, Bryant JA, Sims M, Kraft CS, Cohen SH, Berenson CS, Korman LY, Ford CB, Litcofsky KD, Lombardo M-J, Wortman JR, Wu H, Auniņš JG, McChalicher CWJ, Winkler JA, McGovern BH, Trucksis M, Henn MR, von Moltke L. 2022. SER-109, an Oral Microbiome Therapy for Recurrent *Clostridioides difficile* Infection. N Engl J Med 386:220–229.

20. Sharaby AA, Abugoukh TM, Ahmed W, Ahmed S, Elshaikh AO, Sharaby AA, Abugoukh TM, Ahmed W, Ahmed S, Elshaikh AO. 2022. Do Probiotics Prevent *Clostridium difficile*-Associated Diarrhea? Cureus 14.

21. Allen SJ, Wareham K, Wang D, Bradley C, Hutchings H, Harris W, Dhar A, Brown H, Foden A, Gravenor MB, Mack D. 2013. Lactobacilli and bifidobacteria in the prevention of antibiotic-associated diarrhoea and *Clostridium difficile* diarrhoea in older inpatients (PLACIDE): a randomised, double-blind, placebo-controlled, multicentre trial. The Lancet 382:1249–1257.

22. Gerding DN, Meyer T, Lee C, Cohen SH, Murthy UK, Poirier A, Van Schooneveld TC, Pardi DS, Ramos A, Barron MA, Chen H, Villano S. 2015. Administration of Spores of Nontoxigenic *Clostridium difficile* Strain M3 for Prevention of Recurrent C difficile Infection: A Randomized Clinical Trial. JAMA 313:1719–1727.

23. Leslie JL, Jenior ML, Vendrov KC, Standke AK, Barron MR, O’Brien TJ, Unverdorben L, Thaprawat P, Bergin IL, Schloss PD, Young VB. 2021. Protection from Lethal *Clostridioides difficile* Infection via Intraspecies Competition for Cogerminant. mBio 12:10.1128/mbio.00522-21.

24. Etienne-Mesmin L, Chassaing B, Adekunle O, Mattei LM, Bushman FD, Gewirtz AT. 2018. Toxin-positive *Clostridium difficile* latently infect mouse colonies and protect against highly pathogenic C. difficile. Gut 67:860–871.

25. Wang S, Heuler J, Wickramage I, Sun X. 2022. Genomic and Phenotypic Characterization of the Nontoxigenic *Clostridioides difficile* Strain CCUG37785 and Demonstration of Its Therapeutic Potential for the Prevention of C. difficile Infection. Microbiol Spectr 10:e0178821.

26. Wang S, Zhu D, Sun X. 2022. Development of an Effective Nontoxigenic *Clostridioides difficile*–Based Oral Vaccine against *C. difficile* Infection. Microbiol Spectr 10:e00263–22.

27. Dong Q, Lin H, Allen M-M, Garneau JR, Sia JK, Smith RC, Haro F, McMillen T, Pope RL, Metcalfe C, Burgo V, Woodson C, Dylla N, Kohout C, Sundararajan A, Snitkin ES, Young VB, Fortier L-C, Kamboj M, Pamer EG. 2023. Virulence and genomic diversity among clinical isolates of ST1 (BI/NAP1/027) *Clostridioides difficile*. Cell Rep 42:112861.

28. Carter GP, Lyras D, Allen DL, Mackin KE, Howarth PM, O’Connor JR, Rood JI. 2007. Binary toxin production in *Clostridium difficile* is regulated by CdtR, a LytTR family response regulator. J Bacteriol 189:7290–7301.

29. Lyon SA, Hutton ML, Rood JI, Cheung JK, Lyras D. 2016. CdtR Regulates TcdA and TcdB Production in *Clostridium difficile*. PLoS Pathog 12:e1005758.

30. Labrie SJ, Samson JE, Moineau S. 2010. Bacteriophage resistance mechanisms. Nat Rev Microbiol 8:317–327.

31. van Houte S, Buckling A, Westra ER. 2016. Evolutionary Ecology of Prokaryotic Immune Mechanisms. Microbiol Mol Biol Rev MMBR 80:745–763.

32. Owen SV, Wenner N, Dulberger CL, Rodwell EV, Bowers-Barnard A, Quinones-Olvera N, Rigden DJ, Rubin EJ, Garner EC, Baym M, Hinton JCD. 2021. Prophages encode phage-defense systems with cognate self-immunity. Cell Host Microbe 29:1620–1633.e8.

33. Jarchum I, Liu M, Shi C, Equinda M, Pamer EG. 2012. Critical role for MyD88-mediated neutrophil recruitment during *Clostridium difficile* colitis. Infect Immun 80:2989–2996.

34. Abt MC, Lewis BB, Caballero S, Xiong H, Carter RA, Sušac B, Ling L, Leiner I, Pamer EG. 2015. Innate Immune Defenses Mediated by Two ILC Subsets Are Critical for Protection against Acute *Clostridium difficile* Infection. Cell Host Microbe 18:27–37.

35. Sorg JA, Sonenshein AL. 2008. Bile salts and glycine as cogerminants for *Clostridium difficile* spores. J Bacteriol 190:2505–2512.

36. Battaglioli EJ, Hale VL, Chen J, Jeraldo P, Ruiz-Mojica C, Schmidt BA, Rekdal VM, Till LM, Huq L, Smits SA, Moor WJ, Jones-Hall Y, Smyrk T, Khanna S, Pardi DS, Grover M, Patel R, Chia N, Nelson H, Sonnenburg JL, Farrugia G, Kashyap PC. 2018. *Clostridioides difficile* uses amino acids associated with gut microbial dysbiosis in a subset of patients with diarrhea. Sci Transl Med 10:eaam7019.

37. Bosnjak M, Karpe AV, Van TTH, Kotsanas D, Jenkin GA, Costello SP, Johanesen P, Moore RJ, Beale DJ, Srikhanta YN, Palombo EA, Larcombe S, Lyras D. 2023. Multi-omics analysis of hospital-acquired diarrhoeal patients reveals biomarkers of enterococcal proliferation and *Clostridioides difficile* infection. Nat Commun 14:7737.

38. Karasawa T, Ikoma S, Yamakawa K, Nakamura S. 1995. A defined growth medium for *Clostridium difficile*. Microbiol Read Engl 141 ( Pt 2):371–375.

39. Aguirre AM, Yalcinkaya N, Wu Q, Swennes A, Tessier ME, Roberts P, Miyajima F, Savidge T, Sorg JA. 2021. Bile acid-independent protection against *Clostridioides difficile* infection. PLoS Pathog 17:e1010015.

40. Girinathan BP, DiBenedetto N, Worley JN, Peltier J, Arrieta-Ortiz ML, Immanuel SRC, Lavin R, Delaney ML, Cummins CK, Hoffman M, Luo Y, Gonzalez-Escalona N, Allard M, Onderdonk AB, Gerber GK, Sonenshein AL, Baliga NS, Dupuy B, Bry L. 2021. In vivo commensal control of *Clostridioides difficile* virulence. Cell Host Microbe 29:1693–1708.e7.

41. Hammond GA, Johnson JL. 1995. The toxigenic element of *Clostridium difficile* strain VPI 10463. Microb Pathog 19:203–213.

42. Nagaro KJ, Phillips ST, Cheknis AK, Sambol SP, Zukowski WE, Johnson S, Gerding DN. 2013. Nontoxigenic *Clostridium difficile* Protects Hamsters against Challenge with Historic and Epidemic Strains of Toxigenic BI/NAP1/027 *C. difficile*. Antimicrob Agents Chemother 57:5266–5270.

43. Borriello SP, Barclay FE. 1985. Protection of hamsters against *Clostridium difficile* ileocaecitis by prior colonisation with non-pathogenic strains. J Med Microbiol 19:339–350.

44. Wilson KH, Sheagren JN. 1983. Antagonism of Toxigenic *Clostridium difficile* by Nontoxigenic C. difficile. J Infect Dis 147:733–736.

45. Pruss KM, Enam F, Battaglioli E, DeFeo M, Diaz OR, Higginbottom SK, Fischer CR, Hryckowian AJ, Van Treuren W, Dodd D, Kashyap P, Sonnenburg JL. 2022. Oxidative ornithine metabolism supports non-inflammatory C. difficile colonization.1. Nat Metab 4:19–28.

46. Koenigsknecht MJ, Theriot CM, Bergin IL, Schumacher CA, Schloss PD, Young VB. 2015. Dynamics and Establishment of *Clostridium difficile* Infection in the Murine Gastrointestinal Tract. Infect Immun 83:934–941.

47. Trzilova D, Warren MAH, Gadda NC, Williams CL, Tamayo R. Flagellum and toxin phase variation impacts intestinal colonization and disease development in a mouse model of *Clostridioides difficile* infection. Gut Microbes 14:2038854.

48. Keith JW, Dong Q, Sorbara MT, Becattini S, Sia JK, Gjonbalaj M, Seok R, Leiner IM, Littmann ER, Pamer EG. 2020. Impact of Antibiotic-Resistant Bacteria on Immune Activation and *Clostridioides difficile* Infection in the Mouse Intestine. Infect Immun 88:e00362–19.

49. Fletcher JR, Pike CM, Parsons RJ, Rivera AJ, Foley MH, McLaren MR, Montgomery SA, Theriot CM. 2021. *Clostridioides difficile* exploits toxin-mediated inflammation to alter the host nutritional landscape and exclude competitors from the gut microbiota. Nat Commun 12:462.

50. Oliveira RA, Ng KM, Correia MB, Cabral V, Shi H, Sonnenburg JL, Huang KC, Xavier KB. 2020. *Klebsiella michiganensis* transmission enhances resistance to *Enterobacteriaceae* gut invasion by nutrition competition. Nat Microbiol 5:630– 641.

51. Eberl C, Weiss AS, Jochum LM, Durai Raj AC, Ring D, Hussain S, Herp S, Meng C, Kleigrewe K, Gigl M, Basic M, Stecher B. 2021. *E. coli* enhance colonization resistance against *Salmonella* Typhimurium by competing for galactitol, a context-dependent limiting carbon source. Cell Host Microbe 29:1680–1692.e7.

52. Osbelt L, Wende M, Almási É, Derksen E, Muthukumarasamy U, Lesker TR, Galvez EJC, Pils MC, Schalk E, Chhatwal P, Färber J, Neumann-Schaal M, Fischer T, Schlüter D, Strowig T. 2021. *Klebsiella oxytoca* causes colonization resistance against multidrug-resistant *K. pneumoniae* in the gut via cooperative carbohydrate competition. Cell Host Microbe 29:1663–1679.e7.

53. Rohlfing AE, Eckenroth BE, Forster ER, Kevorkian Y, Donnelly ML, Puebla HB de la, Doublié S, Shen A. 2019. The CspC pseudoprotease regulates germination of *Clostridioides difficile* spores in response to multiple environmental signals. PLOS Genet 15:e1008224.

54. Haak BW, Littmann ER, Chaubard J-L, Pickard AJ, Fontana E, Adhi F, Gyaltshen Y, Ling L, Morjaria SM, Peled JU, van den Brink MR, Geyer AI, Cross JR, Pamer EG, Taur Y. 2018. Impact of gut colonization with butyrate-producing microbiota on respiratory viral infection following allo-HCT. Blood 131:2978–2986.

