## Supplemental figures S1-S5 for "Protection against *Clostridioides difficile* disease by a naturally avirulent *C. difficile* strain"

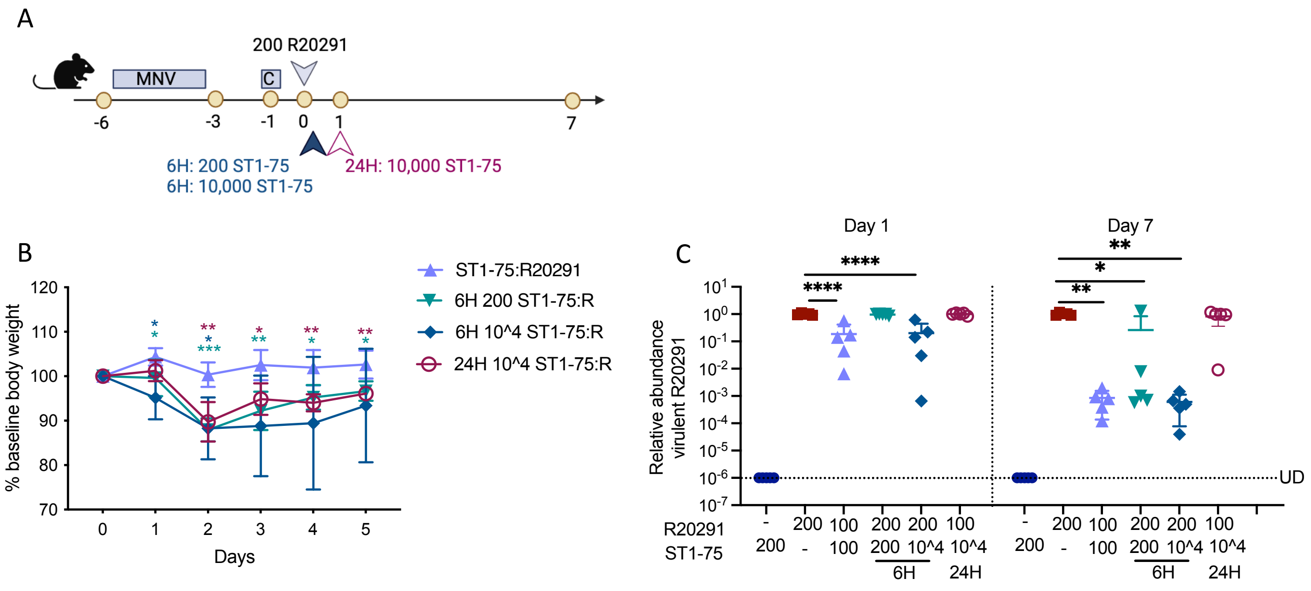


Figure S1 Addition of ST1-75 after inoculation with virulent R20291 fails to protect despite reducing the relative R20291 numbers.

(A) Schematic of the experimental procedure. Wild-type C57BL/6 mice (n = 5 per group) were treated with Metronidazole, Neomycin and Vancomycin (MNV, 0.25 g/L for each) in drinking water for 3 days, followed by one intraperitoneal injection of Clindamycin (200 mg/mouse) 2 days after antibiotic recess. Then, mice were inoculated with 200 R20291 *C. difficile* spores and challenged with ST1-75 *C. difficile* spores via oral gavage at indicated times and doses. Daily body weight was monitored for 5 days post-infection. (B) %Weight loss relative to the baseline of mice infected with indicated strains. Statistical significance was observed between delayed challenge groups against ST1-75:R20291.(C) Relative abundance of R20291 in feces from infected mice was determined by measuring wildtype *cdtR* copies by qPCR on 1 day and 7 days post-infection. UD: Under the limit of detection. Statistical significance was calculated by unpaired t-test and One-way ANOVA, * $p < 0.05$, ** $p < 0.01$, *** $p < 0.001$, **** $p < 0.0001$.

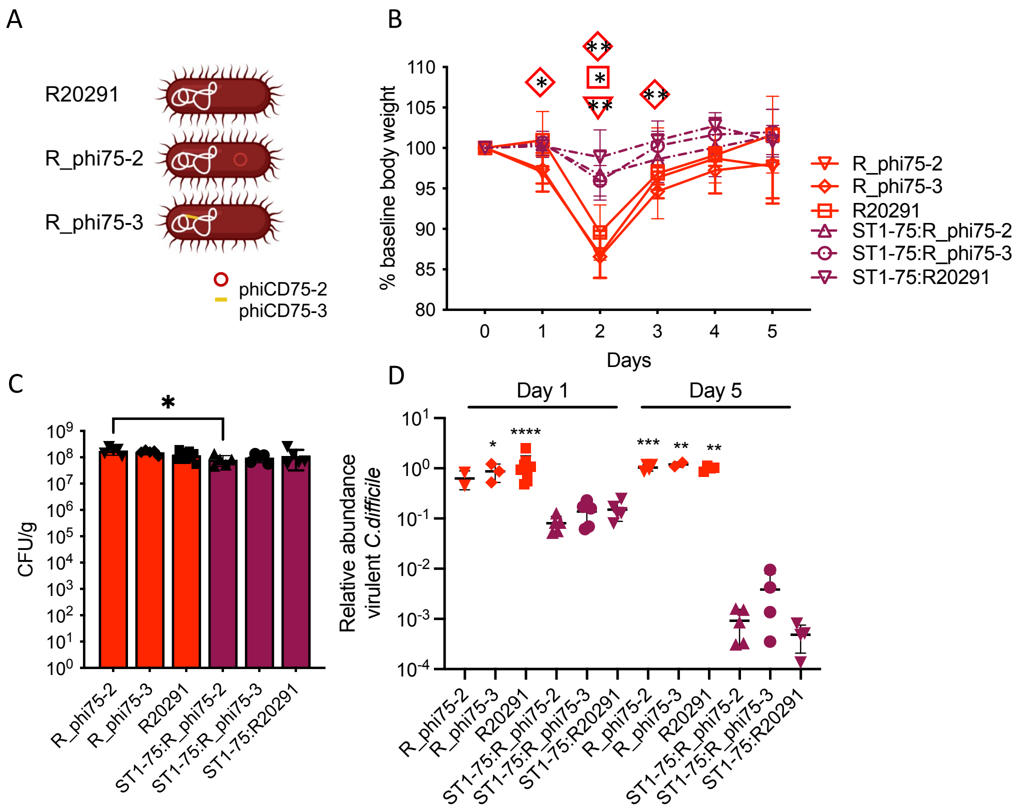


Figure S2. Protection by the avirulent ST1-75 isolate is not due to differences in lysogenic phages between strains.
(A) Lysogeny R20291 strains generated to harbor phiCD75-2 and phiCD75-3, respectively. (B) %Weight loss relative to baseline of mice infected with indicated strains. Statistical significance was observed between R_phi75-2 to ST1-75:R_phi75-2, R_phi75-3 to ST1-75:R_phi75-3, and R20291 to ST1-75:R20291. (C) Fecal colony-forming units were measured by plating on selective agar on 1 day post-infection. (D) Relative abundance of R20291 in feces from infected mice was determined by measuring wildtype *cdtR* copies by qPCR on 1 day and 5 days post-infection. Statistical significance was calculated by One-way ANOVA, * $p < 0.05$, ** $p < 0.01$, *** $p < 0.001$. **** $p < 0.0001$. Statistical significance was observed between R_phi75-2 to ST1-75:R_phi75-2 on day 5, R_phi75-3 to ST1-75:R_phi75-3 on day 1 , and day 5, R20291 to ST1-75:R20291 on day 1 and day 5.

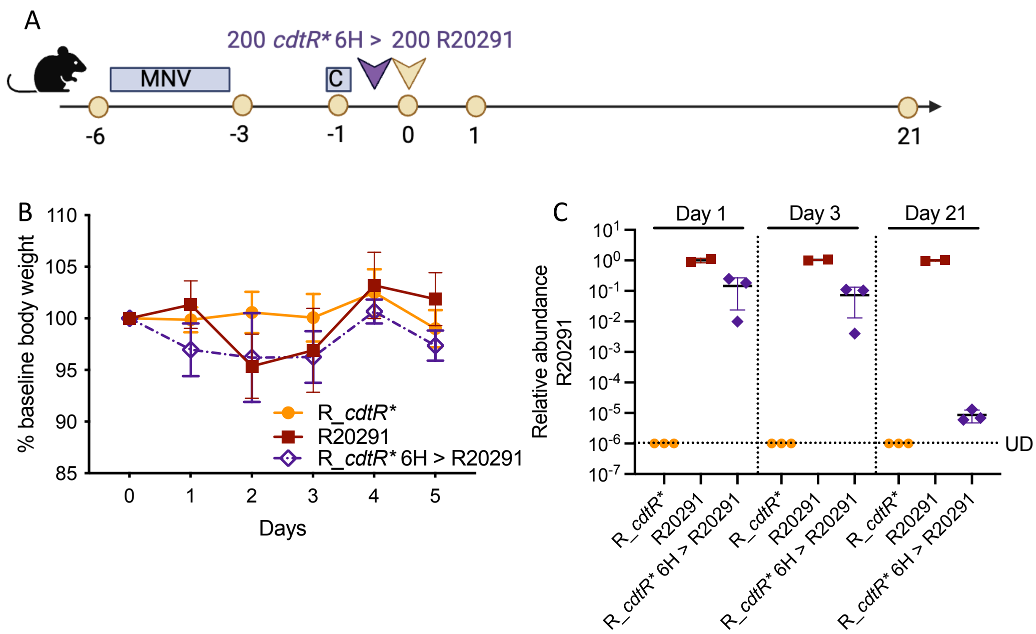


Figure S3 R20291 *cdtR** outcompetes the parental virulent strain in mice.

(A) Schematic of the experimental procedure. Wild-type C57BL/6 mice (n = 2-3 per group) were treated with Metronidazole, Neomycin and Vancomycin (MNV, 0.25 g/L for each) in drinking water for 3 days, followed by one intraperitoneal injection of clindamycin (200 mg/mouse) 2 days after antibiotic recess. Then, mice were inoculated with 200 R_*cdtR** *C. difficile* spores before 200 R20291 *C. difficile* spores via oral gavage. Daily body weight was monitored for 5 days post-infection (B) %Weight loss relative to the baseline of mice infected with indicated strains. (C) Relative abundance of R20291 in feces from infected mice was determined by measuring wildtype *cdtR* copies by qPCR on 1 day, 3 days , and 21 days post-infection. UD: Under the limit of detection.

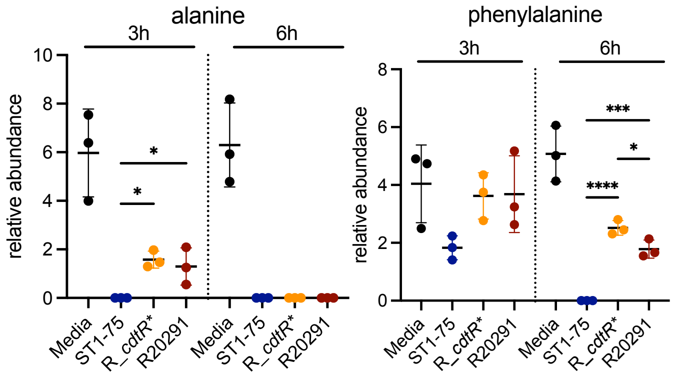


Figure S4 ST1-75 depletes amino acids faster than R_*cdtR** and R20291.
Metabolomic relative abundance of alanine and phenylalanine in cecal extracts inoculated with either ST1-75, R_*cdtR** or R20291.

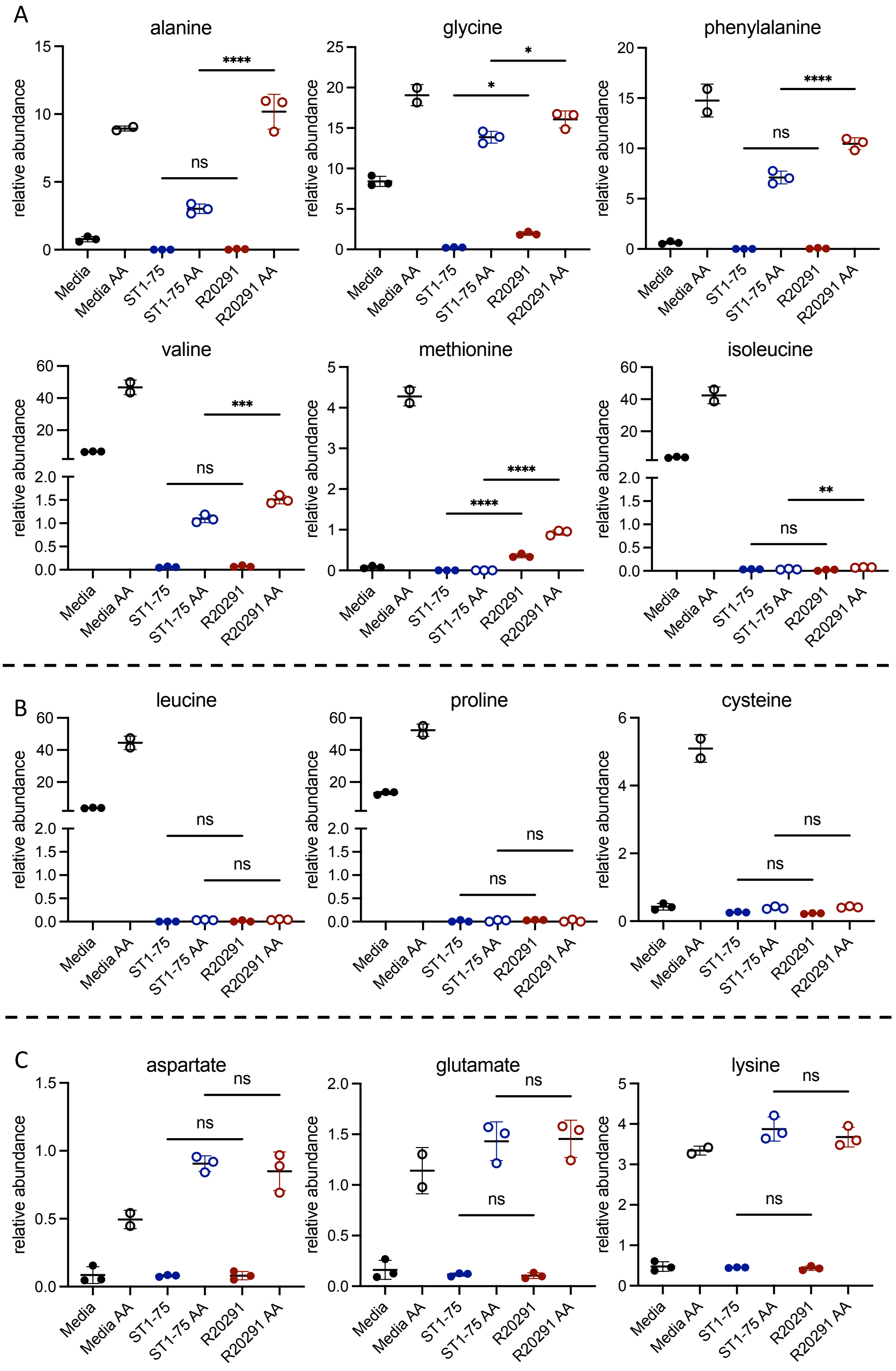


Figure S5 ST1-75 depletes specific amino acids faster than R20291.
Metabolomic relative abundance of each amino acid spiked in cecal extracts and inoculated with either ST1-75 or R20291 for 6 hours. (A) Amino acids depleted faster by ST1-75 than R20291. (B) Amino acids depleted at similar rate by ST1-75 and R20291. (C) Amino acids that were not depleted by ST1-75 or R20291.
